# Does a single exposure to social defeat render rats more vulnerable to chemically induced colitis than brief inescapable foot-shocks?

**DOI:** 10.1101/2022.01.28.478146

**Authors:** Anne Marita Milde, Anne Marie Kinn Rød, Silvia Brekke, Hedda Gjøen, Ghenet Mesfin, Robert Murison

**Affiliations:** Regional Centre for Child and Youth Mental Health and Child Welfare, Norwegian Research Centre, NORCE; Department of Biological and Medical Psychology, University of Bergen, Norway

## Abstract

All social species are to different degrees exposed to stressors being physical or social environmental, which may affect health and well-being. Stressful and traumatic situations have direct effect on immune responses that may underlie susceptibility of developing somatic illness. In animal research, different types of stressors have been investigated in studying the effect on bowel disorders, some stressor being more or less of environmental origin. We aimed therefore to explore whether a more natural stressor would differ from a stressor of more unnatural characteristics on dextran sulphate sodium (DSS) induced colitis in adult rats. Specifically, if social stress within a single social defeat (SD) paradigm would be a more potent stressor than brief inescapable foot-shocks (IFS) in causing elevated faecal granulocyte marker protein (GMP), crypt- and inflammation score in colon tissue. Three groups of male Wistar rats were used; socially defeated rats; inescapable foot-shock rats; and comparison rats. Main findings showed no difference between the groups on GMP levels, however, there was a significant difference on inflammation and crypt score for the distal part of colon, detected through histology, where socially defeated rats were more susceptible. A single SD seems to be more adverse for these animals, but further studies are recommended to validate a broader range of different outcomes comparing two such different rodent stress models.

## Introduction

In humans, at least, the main sources of stress are social, where relationships and sense of belonging are perceived to be threatened. Exposure to social stress varies in magnitude and intensity, and there is a vast amount of evidence for an impact on health [1 for review). Studies have revealed associations between gastrointestinal (GI) diseases such as irritable bowel syndrome (IBS) and inflammatory bowel disease (IBD) and stress, demonstrating the importance of brain-gut interactions [2 for review]. IBD is a collective term for ulcerative colitis (UC) and Crohn’s disease (CD) and differs from IBS, which is a functional GI disorder with a multifaceted pathophysiology. It is unclear whether stressful life events lead to the development of GI diseases, or if they are more strongly related to associated mood disorders. For example, it has been reported that patients with IBD have higher lifetime rates of anxiety and mood disorders, and the onset of these precedes the diagnosis of IBD [3].

In animal research, social defeat (SD) is considered a naturalistic model of social stress. The SD protocol utilizes the resident/intruder paradigm. The paradigm is based on the fact that an adult male rat, the resident, will establish a territory and attack an unfamiliar male, the intruder, when introduced in its home cage. This social conflict situation has numerous effects on behaviour and the neuroendocrine system [4], and a single SD experience can mimic an acute stressor that can provoke long-term fear responses [5]. Inescapable foot-shocks (IFS) are known as a model of acute stress but, unlike SD which relies upon animal-animal interactions, IFS is easily quantifiable and can be precisely controlled. Although it does not normally cause any physical damage, brief exposure to just a few IFS may cause long-lasting behavioural changes [5]. However, critique of using the IFS method includes the pain inflicted on the animal and its artificiality. Less is known of how the more naturalistic stressor (SD) compares to the more artificial one (IFS).

Clinical studies give evidence for significant correlations between increased intestinal permeability and disease activity in ulcerative colitis [6]. Permeability across the colon wall layers allows absorption of nutrients from food and fluids, as well as elimination of waste materials. However, if larger proteins and microorganisms such as bacteria infiltrate these layers, there is a heightened risk of an inflamed colon. Concentrations of faecal calprotectin (FC) are found to be elevated in patients with IBD compared to those with functional GI disorders [7,8,9]. Calprotectin is a protein complex in humans, released extracellularly by activation of neutrophils, and intestinal inflammation causes increased concentrations in the colon [10]. The parallel to human calprotectin, granulocyte marker protein (GMP), can be analysed in rodent faeces [11] thus to detect a disease activity in a non-invasive manner. Here we report both histological data, and measures of GMP in DSS-treated animals after either a single SD or brief IFS.

Previous studies on murine DSS-induced colitis and social stress have utilized models of chronic SD [12]. The present study is to our knowledge the first to assess the outcome of a single social defeat on DSS-induced colitis, and in the same study compare single social defeat to IFS on following chemically induced colon tissue damage in rats. The aim was to compare a single session of social defeat (SD) with those of brief exposure to inescapable foot shocks (IFS) in terms of differences in faecal granulocyte marker protein (GMP) concentrations and colon tissue histology in adult male rats after inducing colitis-like condition by dextran sulphate sodium (DSS). We hypothesized that being physically attacked and defeated would render animals more prone to chemically induced colon tissue damage than would IFS. Further, we explored whether effects would be related to pre-stress levels of corticosterone, as earlier reported for IFS [13].

## Methods and materials

### Animals and housing

All the testing and procedures were approved by the Norwegian Animal Research Authority (permit number: 2006010B) and were registered by the Authority. All effort were made to minimize suffering. The day after arrival, male Wistar rats (9 weeks of age and 260– 299g on arrival) from two separate batches (Taconic, Lille Skvensved, Denmark) were single housed in individually ventilated cages (polypropylene Euro-standard Type III H) with free access to food and water. Within the cages, air was exchanged 75 times per hour, there was an average ambient temperature of 22°C and an average relative humidity of 65%. The room had a 12:12h light/dark schedule with lights on at 07:00h and lights off at 19:00h (progressive increase in light at 06:00h and progressive dimming at 18:00h). Rats were allowed 5 days of acclimatization, then 5 days of daily handling for 1-2 min before blood sampling and experimental procedures.

The resident male rats (Wistar, Taconic) were at least 5 months of age and weighed >450g. To stimulate territorial behaviour, they were housed in pairs together with ovariectomized females (Wistar, Taconic) in individually ventilated cages for at least 2 weeks, housed in a separate colony room, and habituated to being moved. The females were not present during habituation and the SD procedure, but were briefly removed from their cage. For the males, time of transport from the colony room to the test room was approximately 5 min and followed by 1h rest before the introduction of the intruder rat. Bedding was not renewed for at least 2 days prior to a social conflict to preserve the residents’ scent [5].

### Experimental design and procedures

An overview of the experimental design is shown in Fig 1. All male rats (n=90) underwent blood sampling prior to the experimental procedures.

**Figure 1:**
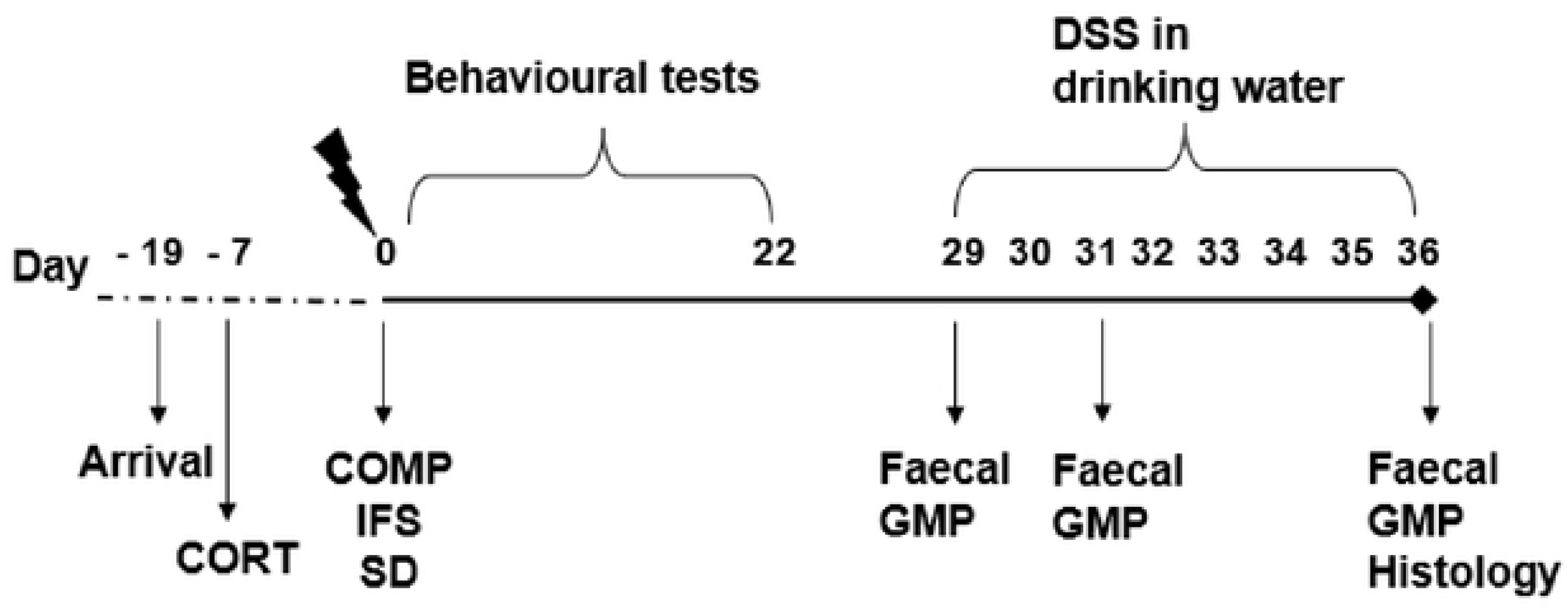
An overview of the experimental design for the three experimental groups: comparison (COMP), inescapable foot shock (IFS) and social defeat (SD) (n=20 pr group). Procedures were identical on all days in all groups except on day O when the different stress procedures were conducted. CORT= blood samples for initial corticosterone measures; Faecal GMP = collection of faecal pellets for measurements of faecal granulocyte marker protein; DSS = exposure to dextran sulphate sodium. Behavioural tests are described in Kinn et al., 2012.

On the basis of the initial corticosterone concentration, the rats were divided into three: high corticosterone (mean ± Standard Error of the Mean (SEM): 239.3 ± 15.8 ng/ml; n = 30), middle corticosterone (106.3 ± 7.8 ng/ml; n = 30) and low corticosterone (32.6 ± 5.1 ng/ml; n = 30). The middle corticosterone group was excluded from the experiment to maximize differences between experimental groups.

From the high and low corticosterone group, rats were divided by stratified randomization into IFS, SD, and comparison (COMP) group (n’s =20). Thus, each of the three experimental groups (IFS, SD and COMP) comprised one high and one low corticosterone subgroup. The subgroups were: IFS-HIGH, IFS-LOW, SD-HIGH. SD-LOW, COMP-HIGH, COMP-LOW (n’s=10).

On day 0 of the experiment each experimental animal underwent IFS, SD or COMP procedures. On all other days, the procedures were identical for all groups. For a separate behavioural experiment ending day 24 (previously reported in [5]), rats were once exposed to acoustic startle response test and once a week for three weeks exposed to sucrose preference test, open field test, elevated plus maze test and body weight were measured.

For chemical induction of colitis, all rats were exposed to 4% DSS in their drinking water from day 29 and for 7 days throughout the experiment.

Faecal pellets for measurements of GMP were collected prior to the DSS exposure, after 2 days exposure to DSS, and after 7 days exposure to DSS. The experimental animals were euthanized day 36 after the stress procedures for histology of colon tissue.

### Corticosterone sampling

To minimize suffering, blood sampling from the saphenous vein was chosen over collection from the jugular vein. One day prior to blood sampling the skin area above the saphenous vein was shaved. Between 09:00 and 12:00 the home cage was moved to the sampling room and the rat placed in a sealed chamber (23×12×11.5 cm) for anesthesia (flow of 1 l/min O_2_, and 1 l/min N_2_O vaporized by 5% isoflurane (Isoba Vet., Schering-Plough, Ballerup, Denmark)). After clear muscle relaxation, the rat was placed in a ventral position, 2% isoflurane was given through a face mask, and the saphenous vein was punctured. 40–400 μl blood was collected in BD Microtainer tubes (Medinor, Oslo, Norway), all within < 3.5 min. Samples were centrifuged at 1600*g* for 10 min, the serum separated and then frozen at −20°C until analysis using Rat Corticosterone Enzyme Immunoassay Kit (DSL-10-81100, MedProbe, Oslo, Norway) with the aid of a plate reader, Wallac 1420 Multilabel counter (PerkinElmer, Oslo, Norway).

### Inescapable foot-shock (IFS) procedures

IFS procedures were performed during the light phase (14:00-18:00h). The shock apparatus consisted of a shock generator (Coulbourn Instruments, Lehigh Valley, PA, USA) and a shock chamber (26×30×30 cm, model H10-11R-TC) with grid flooring inside a sound attenuated cubicle (80×50×50 cm). Foot-shocks were delivered through the grids by a computerized shock system (Habitest system; Graphic State 3.0 software). Each rat was placed individually in the chamber, left undisturbed for 2 min before receiving a total of 10 foot-shocks of 1mA intensity, each of 5s duration, with an inter-shock interval from 24s to 244s (mean 90s). The apparatus was thoroughly cleaned with a 20% ethanol solution between animals.

### Single social defeat (SD) procedures

The single SD procedures were performed between 21:00 and 02:00h (the dark phase). To provide a clear view of the resident–intruder conflict, the room was illuminated by red light, the top lid of the resident’s cage was removed, and an empty cage was placed upside down on top of the cage. The residents had been trained to fight at least five times in confronting younger intruder males (Wistar rats, Taconic). Only residents that had defeated the intruder in < 2 min on the last training session without inflicting injury were selected to proceed. Each SD rat was transported to the experimental room and placed in the cage of the resident. When the SD rat was defeated and showed submissive behaviour (lying motionless on its back), it was moved to a small wire-mesh cage which were in the resident’s cage for a total of 1 hour thus protected from repeated attacks and potential physical injuries.

Immediately after the defeat session, SD rats were returned to their home cages and the colony room.

### Comparison group procedure

These rats were gently handled in the colony room on one occasion for 1 min during the light phase (14:00-18:00h).

### Chemical induction of colitis

All rats were given a 4% solution of dextran sulphate sodium (DSS, powder dissolved in distilled water) (TdB Consultancy AB, Uppsala, Sweden) in place of their normal drinking water. The solution was available *ad libitum* for 7 days, and freshly made each day. Daily consumption was recorded.

### Granulocyte marker protein (GMP)

Fresh faecal pellets were collected from the animal cages

Pellets were stored at −30 °C until analysis. 1g of the sample was diluted in 4 ml extraction buffer (TRIS 12.1g/l, CaCl2 1.47g/l, Mertiolat 0.1g/l dissolved in 0.9% NaCl, pH adjusted to 8.0), and thoroughly homogenized using an Ultra Turrax (2000 r.p.m.) for 20 s or until the material was dissolved. The upper halves of the supernatants were carefully harvested and quantified by GMP ELISA.

### Histology

After 7 days of DSS exposure, the rats were euthanized by CO_2_ followed by dislocation of the cervical vertebrae. The abdominal cavity was opened longitudinally, the colon dislocated from the caecum and the small intestine, flushed with a phosphate buffer in a 10 ml syringe (NaCl, KH2PO4, Na2HPO4×2HH2O), and cut along the mesenteric line. Waste material was carefully removed using the syringe with phosphate buffer. A small segment of the upper and lower colon was discarded due to handling. The remaining distal and proximal segments were cut, gently rinsed, and pinned on a piece of polystyrene with the mucosal layer visible. Each segment was soaked in formalin (4%). Eight sections per segment were stained with hematoxylin and eosin (16 sections per rat), then blindly scored in a randomized manner. Validated scoring systems were used for crypts [14], and inflammation [15]: score 0-intact crypt; score 1-loss of the lower 1/3 of the crypt; score 2-loss of the lower 2/3 of the crypt; score 3-loss of the entire crypt, but intact surface epithelium; score 4-loss of the entire crypt and the surface epithelium. A score of 1 to 4 indicated % surface area affected (1 = 1-25%, 2 = 26–50%, 3 = 51–75%, and 4 = 76–100%). The total crypt score was a product of the crypt score and the affected area score. Inflammation: score 0-normal; score 1-focal inflammatory cell infiltration; score 2-inflammatory cell infiltration, “gland drop-outs” and crypt abscess; score 3-mucosal ulcerations. The score was a product of the inflammation score and the score for the affected area. See Fig 2a for an example of normal colon tissue. Fig 2b demonstrates loss of the entire crypt, loss of the surface epithelium and mucosal ulceration.

**Figure 2:**
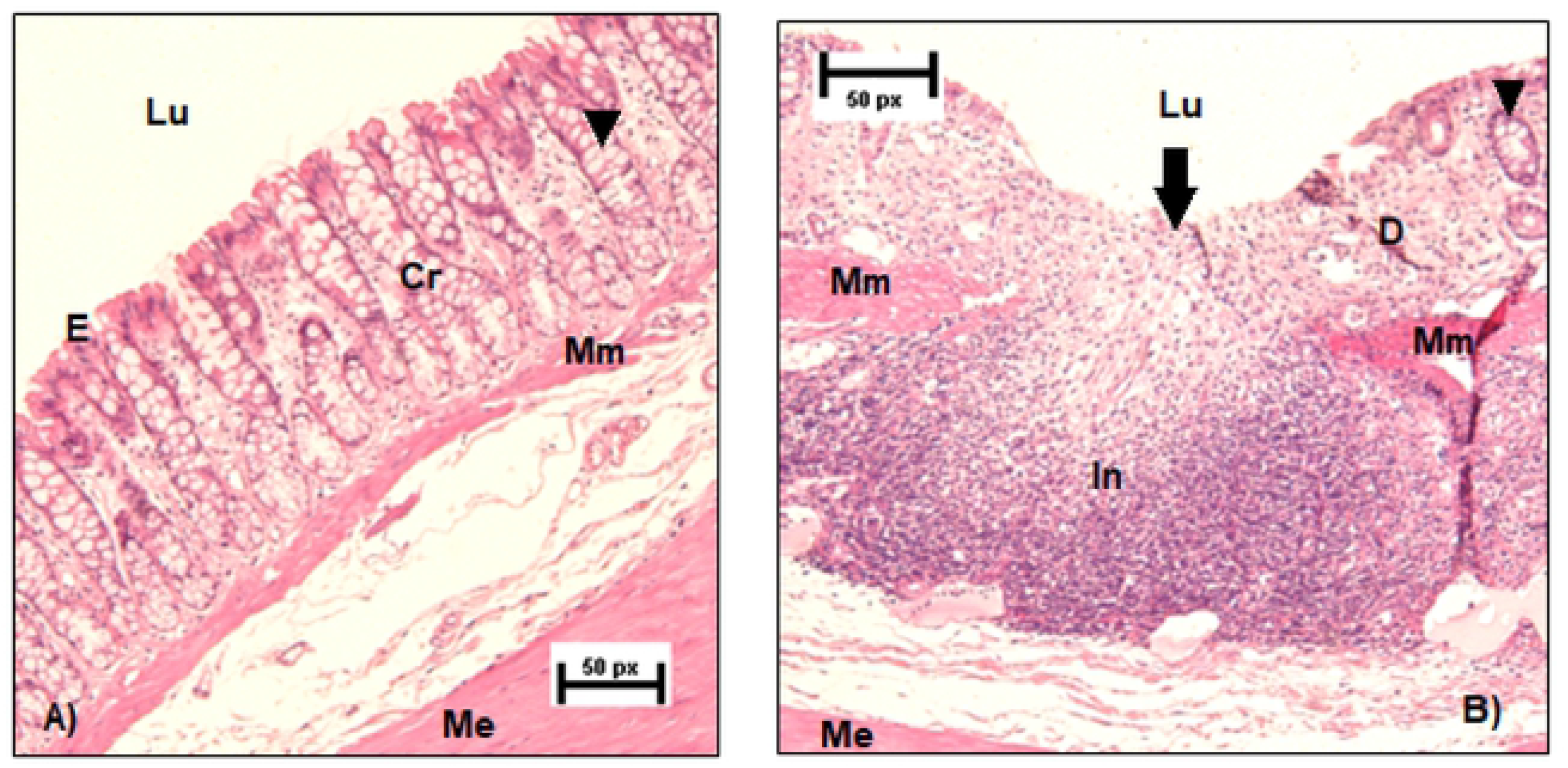
Histology shows Cr (intact crypts), D(degradation), E(surface epithelium), In (infiltrate), Lu (lumen), Mm (muscularis mucosa), Me (muscularis extema), ▾(goblet cells), ↓ infiltrate as degrades crypts and loss of surface epithelium. A) Segment of proximal colon showing intact crypts and no inflammation. B) Segment of distal colon showing colonic damage as crypt losses and infiltration.

The average scores of the eight sections for the proximal segment and for the distal segment on total crypt and total inflammation score were separately compared between the three groups. Interrater agreement was assessed for scoring of 271 histology sections by independent pairs of scorers amongst the five authors in mixed pairs gaining a scoring match of 269 of the sections (interrater agreement of 99.82%).

### Statistics

Statistical analyses were performed using Statistica version 13.3 (TIBCO Software Inc. (2017)). All data are expressed as mean ± SEM. A p-level of <0.05 was considered statistically significant. For total DSS consumption, a factorial ANOVA was used (Group x CORT).

Repeated measures factorial ANOVA was used to analyse the faecal GMP data (group x day x CORT) followed by Bonferroni *post-hoc* analysis. Kruskal-Wallis tests were used to analyse the faecal GMP data between subgroups, due to a significant Levene`s test for (group x day x CORT) in the repeated measures ANOVA analysis. Due to the non-parametric nature of the histology data, Kruskal-Wallis tests were used to compare the three groups, and the six subgroups.

## Results

Three rats were excluded from all analyses. One was excluded from the SD high corticosterone group due to no submission behavior, one from the control high corticosterone group due to technical problems, and one from the IFS high corticosterone group due to overt signs of illness and marked loss of body weight before initiation of procedures for chemically induction of colitis. The included animals were: IFS-HIGH (n=9), IFS-LOW (n=10), SD-HIGH (n=9), SD-LOW (n=10), COMP-HIGH (n=9), COMP-LOW (n=10).

### Consumption of dextran sulphate sodium (DSS) solution

Descriptive statistics of total DSS consumption for experimental groups and subgroups are shown in Table I. Data are expressed as ml.

**Table I.** Faecal GMP levels (mg/1) and Total DSS consumption (ml) Descriptive statistics or faecal GMP levels (mg/I) for comparison group (COMP), inescapab e foot shock group (IFS) and single soc a defeat group (SD), and the groups divided in subgroups according to low and high in tial leve s ofcorticosterone (group mean± SEM). Faecal samples for GMP level ana ys s arc collected prior to dextran sulphate sodium (DSS) exposure (DSS day 0), after 2 days and after 7 days exposure to DSS (DSS day 2 and 7, respect vely). Total DSS consumption (ml) is the total consumption across the 7 days when DSS was g ven in place of the normal drinking water (group mean± SEM). Descriptive statistics of Histology scores from comparison group (COMP) inescapable foot-shock group (IFS) and social defeat group (SD), and the groups div ded in subgroups according to low and high initial levels of corticosterone (group mean± SEM). See text for significant effects and significant differences between groups and between subgroups

| Group | Corticosterone | N | GMP levels<br>Day 0 | GMP levels<br>Day 2 | GMP levels<br>Day 7 | Total DSS<br>consumption |
| --- | --- | --- | --- | --- | --- | --- |
| COMP | Low | 10 | 13.10 ± 2.86 | 49.90 ± 13.81 | 115.18 ± 17.92 | 188.35 ± 8.45 |
| COMP | High | 9 | 22.39 ± 9.48 | 36.00 ± 8.91 | 135.78 ± 21.50 | 197.17 ± 12.19 |
| COMP | All | 19 | 17.50 ± 4.72 | 43.32 ± 8.34 | 124.93 ± 13.70 | 192.53 ± 7.15 |
| IFS | Low | 10 | 12.75 ± 2.23 | 53.48 ± 14.88 | 155.43 ± 40.77 | 188.40 ± 9.57 |
| IFS | High | 9 | 14.22 ± 4.15 | 19.00 ± 6.35 | 93.86 ± 15.82 | 178.33 ± 5.73 |
| IFS | All | 19 | 13.45 ± 2.23 | 37.14 ± 9.12 | 126.26 ± 23.29 | 183.63 ± 5.69 |
| SD | Low | 10 | 19.73 ± 8.12 | 36.20 ± 9.49 | 142.80 ± 34.51 | 192.10 ± 10.48 |
| SD | High | 9 | 8.50 ± 1.34 | 66.31 ± 15.75 | 171.17 ± 40.17 | 186.39 ± 8.65 |
| SD | All | 19 | 14.41 ± 4.41 | 50.46 ± 9.41 | 156.24 ± 25.77 | 189.39 ± 6.71 |

There was no group effect on total DSS consumption, no CORT effect, and no interaction between them (all p’s<0.1).

### Faecal granulocyte marker protein (GMP)

Descriptive statistics of GMP levels (mg/l) across days are shown in Table 1.

Repeated measures factorial ANOVA (group x day x CORT) revealed a significant effect of day, (F(2, 102)=72.51, p<0.001), but no significant effect of either group or CORT (F’s<0.1). None of the interaction effects were significant. Follow-up analysis showed higher levels of GMP after 2 and 7 days of DSS compared to the pre-DSS GMP levels (p=0.02 and p<0.001, respectively). The Kruskal-Wallis test revealed no differences in GMP amongst subgroups on any for the days (all p’s>0.09).

### Histology

Descriptive statistics of histology scores are shown in Table 2.

**Table II.** Histology. Descriptive statistics ofhistologyscores from comparison group (COMP) inescapable fool shock group (IFS) and sing e social defeat group (SD) and 1he groups divided in subgroups according to low and high initial levels ofcorticoslerone (group mean± SEM). Sec text for significant effects and significant differences between groups and between subgroups

| Group | Corticosterone | N | Proximal<br>crypt<br>score | Proximal<br>inflammation<br>score | Distal<br>crypt<br>score | Distal<br>inflammation<br>score |
| --- | --- | --- | --- | --- | --- | --- |
| COMP | Low | 10 | 1.36 ± 0.57 | 0.99 ± 0.42 | 1.98 ± 0.57 | 1.66 ± 0.44 |
| COMP | High | 9 | 1.53 ± 0.53 | 1.18 ± 0.39 | 3.19 ± 0.77 | 2.13 ± 0.46 |
| COMP | All | 19 | 1.44 ± 0.38 | 1.08 ± 0.28 | 2.55 ± 0.48 | 1.88 ± 0.31 |
| IFS | Low | 10 | 1.34 ± 0.53 | 0.90 ± 0.38 | 1.39 ± 0.55 | 0.70 ± 0.37 |
| IFS | High | 9 | 0.42 ± 0.42 | 0.51 ± 0.35 | 1.00 ± 0.55 | 0.56 ± 0.38 |
| IFS | All | 19 | 0.90 ± 0.35 | 0.72 ± 0.26 | 1.20 ± 0.38 | 0.63 ± 0.26 |
| SD | Low | 10 | 0.94 ± 0.48 | 0.81 ± 0.36 | 3.66 ± 0.95 | 2.45 ± 0.51 |
| SD | High | 9 | 1.57 ± 0.58 | 1.29 ± 0.43 | 2.50 ± 0.63 | 2.07 ± 0.52 |
| SD | All | 19 | 1.24 ± 0.37 | 1.04 ± 0.28 | 3.11 ± 0.59 | 2.27 ± 0.36 |

### Proximal colon

For proximal crypt and inflammation scores, there was no significant difference between any of the groups or subgroups (all p’s>0.1).

### Distal colon

For the distal crypt scores, there was a significant difference between experimental groups (H(2, N=57)=7.63, *p* =0.02), where the SD group had higher scores compared to the IFS group (*p*=0.03). There was no significant difference between the IFS and COMP nor between the SD and COMP group, all p’s>0.1. Between the subgroups, there were no significant differences on distal crypt scores (p’s>0.1).

For distal inflammation there was a significant overall group effect (H(2, N=57)=12.87, p<0.002) with higher scores in the SD group compared to the IFS group (p=0.003); further the COMP group had higher scores than the IFS group (p=0.03), but there was no significant differences between the subgroups, all p’s>0.1.

## Discussion

The aim of the current study was to examine whether a single social defeat exposure would differ from brief inescapable foot-shocks in its impact on experimentally induced colonic inflammation in rats. Specifically, whether single SD would cause greater elevations of faecal GMP and render the colon tissue more susceptible to damage than brief IFS. The results confirm previous findings that unrestricted oral ingestion of DSS is an effective method for inducing a colitis-like condition in rodents [16,17]. We did not find a significant overall difference between groups on GMP levels. Exposure to single SD in general had significant impact on histological measurements (crypt and inflammation scores) in the distal colon. Effects were not related to pre-stress levels of corticosterone. There are some additional findings that seem challenging to interpret, such as higher inflammation scores in animals not prior exposed to a stressor compared to animals exposed to foot-shocks.

Animal models of inflammatory bowel disease (IBD) are of value due to their wide range of options for investigating the various factors related to pathogenesis and to develop and evaluate medical treatment options. However, induced colitis models do not reproduce the complexity of the disease, and there are some limitations considering diagnostic subcategories that are rarely investigated in animals.

A valid model of human UC requires that animals develop a colitis-like condition with prominent neutrophils in the epithelium, cryptitis, crypt abscesses and erosions. High faecal levels of calprotectin indicate intestinal inflammation, and due to the simple and non-invasive sampling, numerous tests can be performed repeatedly on the same individual. We collected faeces from the rats thus enabling detection of the rodent parallel to calprotectin, granulocyte marker protein (GMP), where high levels are strong indicators of colonic inflammation [18,19]. Overall, longer exposure to DSS caused higher GMP levels. When analysing levels of GMP prior to the administration of DSS, after 48 hours and after 7 days on DSS, we were unable to find significant differences between rats exposed to Single SD, IFS, and those not prior exposed to an intended stressor. Thus, neither of the two stress procedures rendered the animals more sensitive to DSS as measured by GMP.

The histology data in the present study revealed that most of the erosions occurred in the distal colon and not in the proximal part of the colon. This result is in line with clinical findings were some patients with proctitis or left-sided colitis might also have a caecal patch of inflammation. Bloody diarrhoea is the characteristic symptom of the disease, but supportive findings are vital for establishing the diagnose [20 for update].

Importantly, the anatomy of rodent intestines is unlike human anatomy were the left sided colitis is parallel to the subdivided distal part. In rats, the distal part lays on the right side where the muscular layer is thicker than the proximal left colon. Nonetheless, there is evidence for distinguishing subdivisions of the rat colon thus resembling human anatomy [21]. In our study, rats exposed to Single SD had significantly higher crypt scores and inflammation scores in the distal part of the colon compared to rats exposed to IFS which may indicate differences in the consequence of being exposed to the more natural stressor of physical attack versus an “unnatural” stressor on immune function.

Social defeat, as a result of territorial aggression is associated with emotional stress. It would be of interest to further study differences in aggressive behaviour in relation to susceptibility to induced intestinal inflammation. Studies have shown that animals with an aggressive and proactive coping style (fighting back during a SD confrontation) tend to have a higher sympathetic stress reactivity and lower corticosterone response than do reactive coping animals (non-aggressive submissive) which have a higher parasympathetic response and in general react with the highest corticosterone response [22]. Strategies of stress coping in humans are related to differences in immuno-stimulation [23]. Investigation of stress coping may therefore be of importance in animal studies on ulcerative colitis when using DSS after single SD or even other experimental stressors.

Surprisingly, rats that had not been exposed to either IFS or single SD had significantly higher inflammation scores in the distal colon compared to those who had received foot-shocks. We have no reason to underestimate the effects of IFS, but further studies should investigate the time course of recovery from various stressors. The effect of ten foot-shocks might have diminished to such an extent that these animals were more comparable to those left undisturbed.

Most pre-clinical models on stress-related disorders have focused on male rats, despite that woman seem to be more susceptible to stress-related symptoms in general. Here, we used only male rats, as the social defeat model was chosen as the social stressor, and the same gender should be used for subjects of both single SD and IFS. Studies on social defeat have traditionally been restricted to males since this model relies on territorial aggression between male rodents. This innate behavior is not normally seen in female rodents [24]. In addition, male rodents in general does not display aggressiveness towards females when introduced into their territory if the male resident is not a pathological aggressor [25]. In later years, efforts have been made to establish social stress models using female rats [24]. Future studies on social stress models should consider using female rodents. In humans, the incidence of ulcerative colitis is similar in men and women before the age of 45, whilst above the age of 45, men have a higher risk than women [26]. It is therefore recommended to include both males and females if feasible when modelling human colitis. We are aware that to strengthen the validity of the present study, several immune outcome measures should be included to better establish the presence of inflammation such as the concentration of pro-inflammatory cytokines in mucosal tissue. Further, this model in combination with stress-exposure, is suitable to demonstrate prevention and treatment of induced colitis.

In conclusion, rats previously exposed to a single SD had more colonic damage compared to rats exposed to brief IFS, were both groups had undergone similar experimentally inducement of colitis. The Single SD in general had significantly more impact on histological measurements than brief IFS, shown by higher crypt and inflammation scores in the distal colon, but no difference in GMP measurements was found between the groups. The higher grade of inflammation and tissue damage in the distal part of the colon in socially defeated rats may indicate presentation of differences between the outcomes of ‘natural’ versus ‘unnatural’ stressors. A single social defeat seems to be more adverse for the animals in the present study, but further investigations are recommended to validate a broader range of different outcomes comparing two such different rodent stress models in addition to the aspect of recovery.

## Acknowledgements

This study was performed with support from the Faculty of Psychology, University of Bergen, Norway

